# MechanoAge, a machine learning platform to identify individuals susceptible to breast cancer based on mechanical properties of single cells

**DOI:** 10.1101/2025.08.08.668946

**Authors:** Stefan Hinz, Sturla M. Grøndal, Masaru Miyano, Jennifer C. Lopez, Kristen L. Cotner, Taylor Thomsen, Chang Chen, Edward J. Hester, Lisa D. Yee, Victoria E. Seewaldt, James B. Lorens, Lydia L. Sohn, Mark A. LaBarge

**Author notes:** Corresponding authors:* Mark LaBarge,; Stefan Hinz.

## Abstract

**Background:** Existing breast cancer risk models inadequately identify individuals at latent risk, particularly among women without known genetic mutations or family history. Risk is often underestimated or overestimated due to reliance on population-level data and neglect of cellular aging and mechanobiological alterations.

**Methods:** We profiled primary human mammary epithelial cells (HMECs) from women of varying ages and risk backgrounds using mechano-node-pore sensing (mechano-NPS), a high-throughput microfluidic platform that captures single-cell mechanical properties. Using machine learning, we developed a classifier, MechanoAge, to predict age-related mechanical phenotypes and introduce a novel index, mechano-RISQ, to quantify deviations linked to breast cancer risk. We further assessed the cytoskeletal protein keratin 14 (KRT14) as a molecular mediator of these mechanical states through overexpression and knockdown experiments.

**Findings:** Cells from younger women carrying BRCA1/2 mutations or with a family history of breast cancer exhibited accelerated mechanical aging compared to age-matched controls. Elevated mechano-RISQ scores reflected an increased proportion of cells with “older” mechanical profiles. KRT14 overexpression induced an aged mechanical phenotype in younger cells, while knockdown partially reversed this state in older cells. CyTOF profiling and modeling showed KRT14 modulation impacted protein expression signatures associated with aging and risk, particularly in luminal cells.

**Interpretation:** Mechanical properties of breast epithelial cells reflect biologic aging and cancer susceptibility. Mechano-RISQ offers a new approach for identifying individuals at elevated risk, especially among average-risk populations, and may complement existing risk models by incorporating biophysical measures of epithelial aging.

## Introduction

Breast cancer, as one of the most frequently diagnosed cancers worldwide and a leading cause of cancer-related mortality among women, has long been the subject of efforts to improve risk stratification and early detection strategies. Despite considerable advances in both screening technologies and therapeutic interventions, accurately determining which individuals, particularly among those considered average-risk, are most likely to develop the disease remains one of the most persistent challenges in oncology and public health.

Currently, risk models such as the Gail and Tyrer-Cuzick attempt to integrate clinical and demographic variables including age, reproductive history, family history, and benign breast disease to estimate breast cancer risk over defined time periods^1^. These models, although widely used in clinical practice, suffer from known limitations: the Gail model tends to underestimate risk in many women, while the Tyrer-Cuzick model has been shown to overestimate risk, especially in those with atypical hyperplasia on biopsy^2^. More importantly, both models rely on population-level data and offer limited insight into an individual’s ongoing biological responses to aging or environmental exposures. In particular, they fall short in accounting for how biological processes in breast cells affect an individual’s underlying susceptibility to the disease.

Genetic testing for high-penetrance mutations such as those in BRCA1 and BRCA2 has yielded powerful predictive tools for the small subset of women whose risk is hereditary, which are estimated to comprise just 5–10% of all cases. Women harboring these mutations may face a lifetime breast cancer risk as high as 83%^3,4^, and first-degree family history confers a lifetime risk of approximately 21%^5^. Yet a majority of breast cancers arise in women with no family history and no known pathogenic variants, whose estimated lifetime risk is 13%. Among this large group of ostensibly average-risk individuals, it remains difficult to identify those with latent risk that stems from cellular, molecular, and biophysical alterations that current models are not designed to capture.

This gap is especially concerning in the context of early-onset breast cancer (EOBC), where incidence has been rising at an alarming rate. Between 2012 and 2021, breast cancer incidence increased by 1.4% annually in women under 50, compared to 0.7% in older women, and was nearly twice that rate in certain ancestry groups^6^. These trends suggest that ancestry-related biology and non-hereditary contributors play a larger role than previously appreciated. Environmental exposures, lifestyle changes, delayed childbearing, and rising prevalence of obesity are all potential contributing factors, yet we still lack a predictive framework that incorporates how these external variables interact with internal biological aging processes in the mammary epithelium.

To address this need, we propose that changes in the mechanical properties of cells, which are long recognized as hallmarks of both aging and disease^7^, offer a previously unexamined opportunity to assess individual susceptibility to breast cancer. Aging and disease progression are associated with alterations in cellular elasticity, stiffness, and morphology, which stem from cytoskeletal remodeling, matrix interactions, and disrupted mechanotransduction pathways. These mechanical changes are functional reflections of the underlying cell state and also contain discriminative information capable of resolving subtle differences in lineage commitment, chronological age, and malignant potential.

In previous work, we showed that genome- and proteome-wide changes in human mammary epithelial cells (HMECs) with age correlate with high-risk phenotypes, and that the cytoskeletal intermediate filament protein keratin 14 (KRT14) is differentially expressed in older mammary luminal epithelial cells^8^. Crucially, hallmarks that we associated with aging including this key cytoskeletal change, are accelerated by decades in young women who carry BRCA1, BRCA2, or PALB2 germline mutations compared to average risk^9,10^. These observations suggest that biophysical phenotyping may serve as a surrogate for biological aging and cancer risk, particularly in epithelial lineages of the breast. To quantify these properties at single-cell resolution and in high throughput, we employed mechano-node-pore sensing (mechano-NPS), a microfluidic platform that yields large, multi-parametric datasets capturing mechanical features of individual cells^11,12^.

Here, using mechano-NPS, we mechanically profiled primary HMECs from donors of varying ages and cancer risk levels and applied machine learning to classify mechanical phenotypes as a function of age in average risk women. These phenotypes were predictive of breast cancer susceptibility and associated with accelerated mechanical aging in young women with germline mutations in BRCA1 or BRCA2, or with strong family histories of breast cancer. We introduce a new index, mechano-Risk Index for Single-cell Quantification (mechano-RISQ), to describe this biophysical signature of cancer-prone aging epithelia. Furthermore, we investigated the role of KRT14 in modulating this phenotype and found it to be integral to the mechanical state of high-risk cells.

While other aging biomarkers such as epigenetic clocks or telomere length provide valuable perspectives, they capture only slices of the complex and multilayered biology of aging. By contrast, mechanical phenotyping, anchored in cytoskeletal dynamics and cell-matrix interactions, represents an integrative, systems-level perspective that directly reflects cellular function and risk state. As such, we propose mechano-RISQ as a novel class of biomarker that fills a longstanding void in risk prediction: identifying individuals with accelerated aging-driven susceptibility to breast cancer before malignancy emerges.

## Results

### Database Creation

We used primary HMECs from 18 women of varying ages (**Supplemental Table 1**) as a model system to investigate aging-related changes (**Figure 1A**). HMECs offer a unique advantage because they can be obtained from multiple donors, allowing the analysis of a diverse range of specimens. Unlike immortalized cell lines, primary HMECs maintain genomic stability and acquire minimal tissue culture adaptations, providing a biologically relevant system for examining age-associated cellular changes^13,14^. HMECs are neither cancerous nor immortalized, enabling the exploration of aging processes by comparing cells obtained from women of different chronological ages.

**Figure 1.**
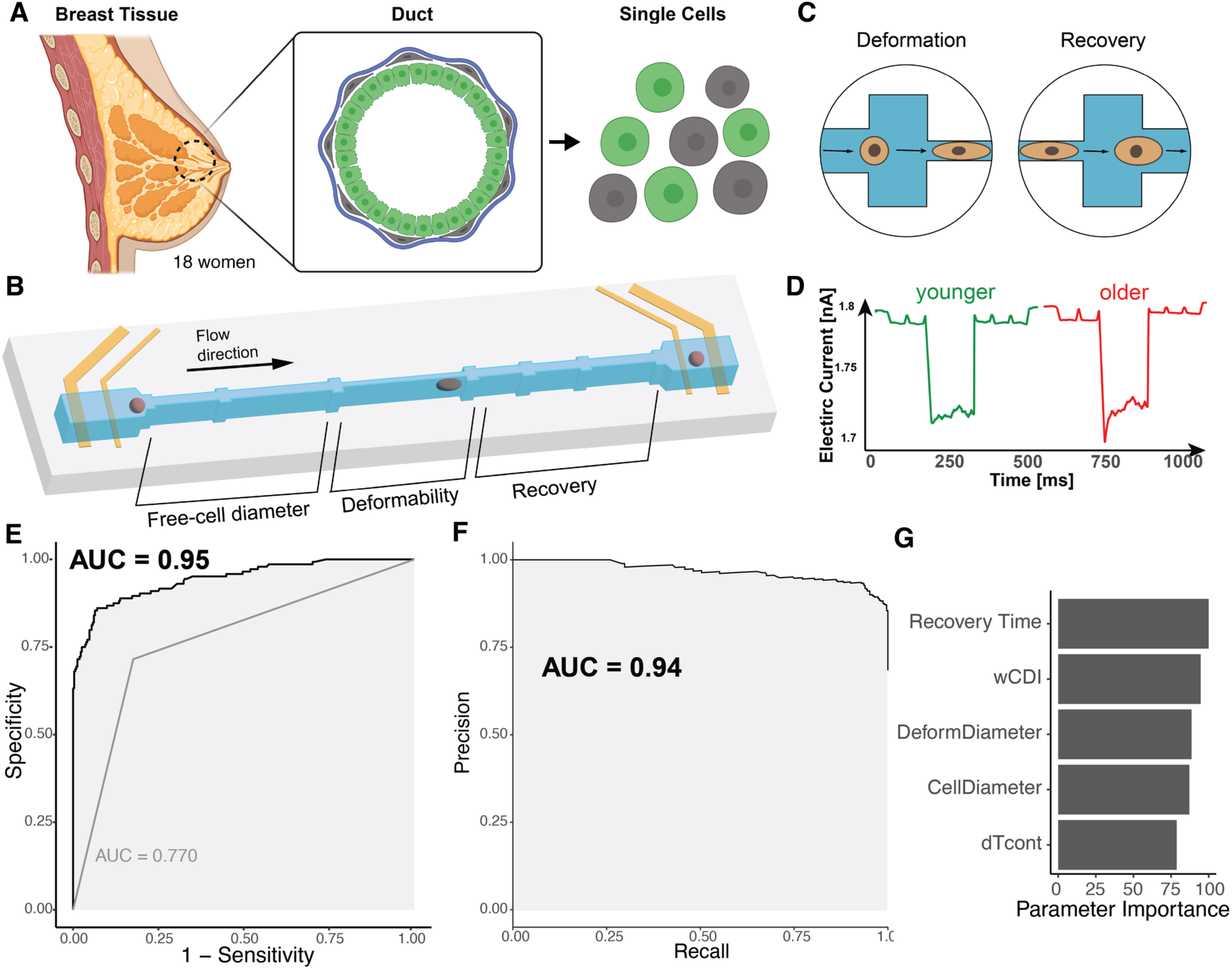
Age-Dependent Mechanobiological Profiles in Breast Cancer. (A) Schematic of breast tissue structure and cell isolation from 18 women. (B) Overview of the mechano-NPS microfluidic device used to measure single-cell mechanical properties. (C) Example cell deformation and recovery process within the mechano-NPS device. (D) Representative pulses of electric current, illustrating current drops associated with cell deformation and recovery within the microfluidic channel. (E) ROC curve illustrating the performance of MechanoAge’s classification of age groups based on validation data. (F) Precision-recall curve of the mechanobiological classifier in predicting age. (G) Feature importance derived from the mechanobiological classifier, highlighting the top five parameters contributing to the risk prediction.

We measured the mechanical properties of the cells via mechano-NPS. This platform consists of a microfluidic channel segmented by larger areas called “nodes”; one segment between two nodes, the “contraction channel”, is narrower than the diameter of a cell, causing the cell to experience a constant strain of *ε* = 0.3 in these studies for a defined period (**Figure 1B** and **1C**). A four-terminal measurement of the current across the microfluidic channel measured the modulating current caused by a transiting cell. An analysis of the modulated current provided information on cell diameter, stiffness, and recovery from deformation (**Figure 1D**, see Methods). By leveraging HMECs in conjunction with high-resolution mechanical data, we quantified mechanical changes across a broad age spectrum. We created a training dataset comprising over 1500 single-cell mechanical profiles from 18 women.

### Model Performance

We modeled the relationship between cellular mechanical features and age using ensemble machine learning algorithms, including Bagged Decision Trees, Random Forests, and Extremely Randomized Trees. These algorithms are well-suited for biological data as they effectively capture non-linear interactions, handle high-dimensional datasets, and are robust against noise, predictor correlations, and the variability inherent in biological measurements^15^. Our model — MechanoAge — predicts the age category of HMECs at the single-cell level based on mechanical phenotypes measured using the mechano-NPS platform. MechanoAge has a high predictive accuracy: the Receiver Operating Characteristic (ROC) curve, with an area under the curve (AUC) of 0.95 (**Figure 1E**), indicates the MechanoAge’s exceptional ability to distinguish between different age categories of HMECs. This performance assessment was based solely on validation data that was not used during training. The ROC curve indicated a strong balance between sensitivity and specificity across various thresholds and ensured accurate classification of HMECs into younger or older categories. This robustness across thresholds suggested that the MechanoAge consistently predicted well, across a wide range of cut-off points used. The Precision-Recall (PR) curve yielded an AUC of 0.94 (**Figure 1F**), which further validated the MechanoAge’s reliability on validation data. The PR curve’s high AUC confirmed the MechanoAge’s resilience to class imbalances and achieved high precision without significantly increased false positives. Thus, MechanoAge was sensitive to true mechanobiological age-related signals and remained resistant to variations in the test data. **Figure 1G** illustrates the importance of the top five mechanical properties measured by mechano-NPS platform in predicting the age of HMECs. “Recovery Time” emerged as the most critical feature and indicated that the time it took for a cell to recover after deformation was a key determinant in classifying its age category. For training purposes, recovery time was divided into five distinct binary variables; the cumulative contribution of these variables is shown **Figure 1G**. This was closely followed by “wCDI”, a dimensionless parameter defined by Kim *et al.* that was inversely related to stiffness, “Deform Diameter” (the diameter of a cell when it is in the contraction channel in the mechano-NPS), and “Cell Diameter” (the “free” diameter of the cells), all of which significantly enhanced the MechanoAge’s predictive power *(3)*. A lesser yet still significant importance was attributed to “ΔTcont” (the cell’s transit time through the contraction channel).

When only considering the fraction of slow-recovering cells after deformation, as previously observed by Kim et al. *(3)*, the property showed a striking decrease in performance, with an AUC of 0.77 (**Figure 1E**). Whereas Recovery Time was a significant individual predictor, the lower AUC compared to that from MechanoAge indicated that this parameter alone, did not capture the full complexity of the data. The reduced AUC signified a diminished capability to accurately classify cells into their corresponding age categories because the additional mechanical characteristics provided complementary information. Thus, implying that the various mechanical properties measured by the mechano-NPS platform interacted in critical ways for differentiating between age categories and that relying on a single feature led to a significant loss of information.

### Mechanical Age as a Predictor of Risk

Age is the principal risk factor for developing breast cancer^16^. We previously showed that some biochemical and transcriptomic breast-specific aging hallmarks were accelerated in women who harbored pathogenic alleles of BRCA1, BRCA2, or PALB2^17,18^. Given the ability of our MechanoAge to accurately predict the age category based on mechanical characteristics, we determined whether mechano-NPS could detect aberrations in “mechano-age”, which would then serve as early indicators of elevated breast cancer risk in women with high-risk mutations.

We applied the MechanoAge to HMECs from women with pathogenic alleles of BRCA1 and BRCA2 or with a family history of breast cancer but without any recognized high-risk alleles (**Figure 2**). Astoundingly, a significant proportion of younger BRCA2 carriers (HR-1 and HR-3) and BRCA1 carriers (HR-2) cells (47%, 79%, and 61%, respectively) were classified as “older” (**Figure 2A**). The age predictions based on mechanobiological properties shifted towards an older phenotype in high-risk mutation carriers. Accelerated aging signals in these high-risk groups manifested as emergent mechanical properties, which emphasizes the potential for early detection of cellular changes in individuals with elevated breast cancer risk.

**Figure 2:**
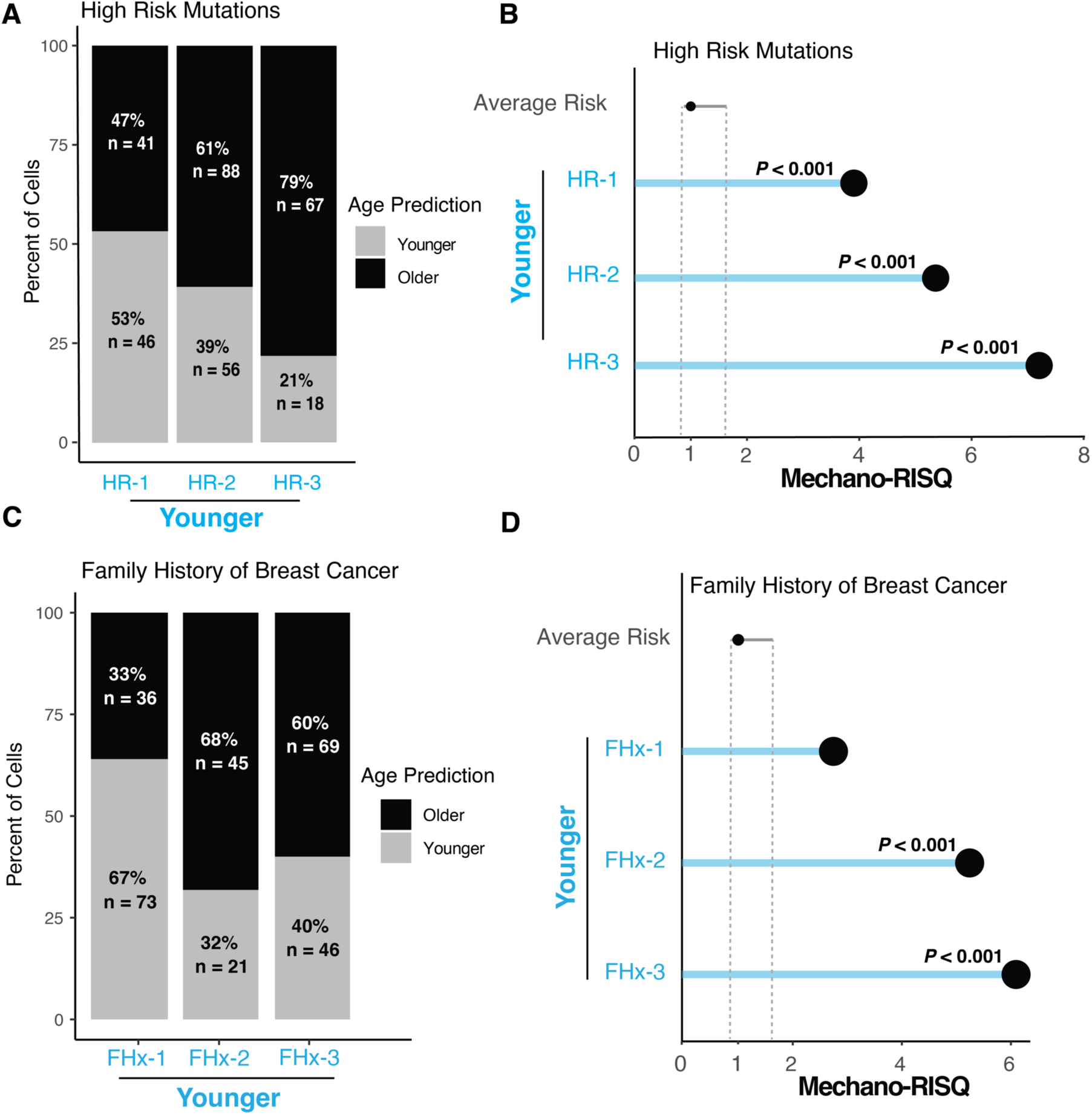
Age-Linked Mechanobiological Profiles Enhance Risk Stratification in Breast Cancer. (A) Bar plots showing the percentage of cell age classification based on their mechanobiological signatures in high-risk (HR) patients. (B) Dot plot illustrating the Mechano-RISQ in individuals with high-risk mutations. Significance was determined using Crawford-Howell test. Dotted line: 95% confidence interval. (C) Stacked bar plots showing the percentage of cell age classification in individuals with a family history of breast cancer. (D) Dot plot demonstrating the Mechano-RISQ for individuals with a family history of breast cancer. Significance was determined using Crawford-Howell test. Dotted line: 95% confidence interval.

To quantify our age-based prediction, we developed the Mechano-RISQ score, an odds ratio-inspired metric that compared the expected average risk misclassification to the observed classification error. Mechano-RISQ scores for HR samples were elevated compared to those of average-risk individuals (**Figure 2B**). Specifically, cells isolated from younger HR women showed significantly higher Mechano-RISQ scores (*P* < 0.001 for HR-1, HR-2, and HR-3). This pattern parallels our previous findings in with an orthogonal molecular assay that found germline high-risk mutations similarly advance biological age estimates derived from an epigenomic ELF5 clock, reinforcing the concept that hereditary risk accelerates aging metrics of mammary epithelium.

We next evaluated cells from women with a family history (FHx) of breast cancer without any identified HR mutations (**Figure 2C**), a group for whom the assessment of true risk was more challenging due to the absence of definitive genetic markers. This group of younger FHx individuals exhibited a higher proportion of cells classified as “older” than expected. The cells isolated from younger FHx individuals exhibited significantly elevated Mechano-RISQ scores compared to the average-risk population (*P* <0.001 for FHx-2, FHx-3), which indicated an aged mechanical phenotype (**Figure 2D)**. Thus, cells from these individuals with a family history of breast cancer, even in the absence of detected known high-risk alleles, had emergent mechanical properties shared with cells known to be HR.

Our cellular mechanical classifier highlighted differences in age prediction accuracy among pathogenic BRCA1/2 mutation carriers and those with a family history of breast cancer. Their elevated Mechano-RISQ scores suggested that these cellular aging profiles deviated from average risk patterns and potentially reflect underlying cancer susceptibility.

### Keratin 14 as a Molecular Mediator of Age-Associated Mechanical States

To investigate the molecular underpinnings of the observed mechanical changes and their effect on age predictions, we focused on KRT14, an intermediate filament and marker of myoepithelial cells in younger mammary epithelia. KRT14 upregulation occurs with age and in genetic high risk in luminal cells^8,19,20^. Given KRT14’s role in cytoskeletal organization, we explored how its overexpression and knockdown influenced mechanobiological properties, specifically regarding age prediction accuracy. We examined the effects of KRT14 overexpression (KRT14 OE) in luminal HMECs from three different average-risk (AR) younger women (AR-1, AR-2, and AR-3). KRT14 OE increased the proportion of cells classified as older in all three strains, with significant differences in AR2 and AR3 (*P* < 0.05, Fisher’s exact test), and AR1 displayed a similar trend (**Figure 3A**). KRT14 overexpression in the HMECs resulted in significantly elevated Mechano-RISQ scores (*P* < 0.05 and *P* < 0.01, respectively), which indicated that KRT14 increased the probability of younger cells being classified as older (**Figure 3B**).

**Figure 3.**
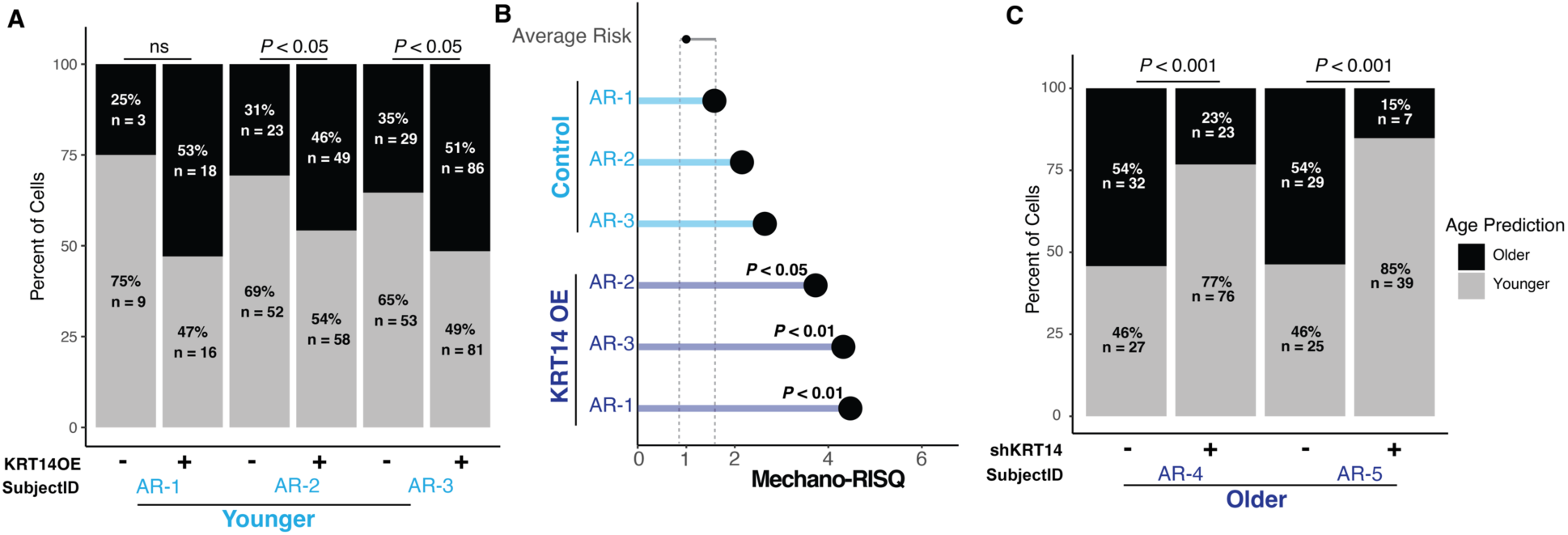
Modulation of Keratin 14 Alters Predicted Cellular Age and Mechano-Risk Score. (A) Bar plots showing the percentage of cells predicted as older or younger based on the age prediction MechanoAge across KRT14 overexpression (KRT14OE) and control conditions in luminal cells. Significance was determined using Fisher’s exact test. (B) Dot plot showing Mechano-RISQ scores for younger cells under KRT14OE and control conditions in luminal cells. Statistical significance is shown for changes in risk scores. Significance was determined using the Crawford-Howell test. Dotted line: 95% confidence interval. (C) Bar plots showing the percentage of cells predicted as older or younger across shKRT14 and control conditions in older luminal cells. Significance was determined using Fisher’s exact test.

Next, we examined the effects of KRT14 knockdown (shKRT14) in older luminal cells and focused on two individuals (AR4 and AR5). KRT14 knockdown led to a significant reduction in the proportion of cells classified as “older” in both (*P* < 0.001) **(Figure 3C)**. In AR5, only 15% of cells were classified as older following KRT14 knockdown, compared to 85% classified as younger. A similar distribution was observed in AR4, where the proportion of cells classified as older dropped to 23%, which indicated that KRT14 knockdown in older cells substantially reduced older mechanical phenotype. This suggested a critical role of KRT14 in the mechanical aging process, and that its knockdown partially reversed the age-related mechanobiological changes in older cells.

Results demonstrated that KRT14, a key cytoskeletal protein, significantly influenced the mechanical properties of younger and older cells. Overexpression of KRT14 in younger cells promoted an “older” mechanical phenotype, whereas the knock down of KRT14 in older cells reduced their likelihood of being classified as older. This highlights KRT14’s central role in age-associated mechanical states.

### KRT14 Modulates Molecular Signatures and Mechanical Phenotypes

To investigate age- and risk-associated protein expression associated with mechanical states, we analyzed ∼14,000 HMECs from three younger AR, three older AR, and six HR individuals. Using CyTOF, we assessed 27 proteins and phospho-proteins relevant to cell-cycle regulation, signaling, and cytoskeletal dynamics (**Supplemental Table 2**). The panel was adapted from a previously validated framework (Pelissier-Vatter *et al*. *(9)*), augmented with additional markers implicated in cytoskeletal regulation. UMAP visualization of control HMECs (**Figure 4A**) showed distinct aggregation of luminal and myoepithelial populations, with notable variability in protein expression for known lineage markers such as KRT7, KRT19, CD271, and CD44.

**Figure 4.**
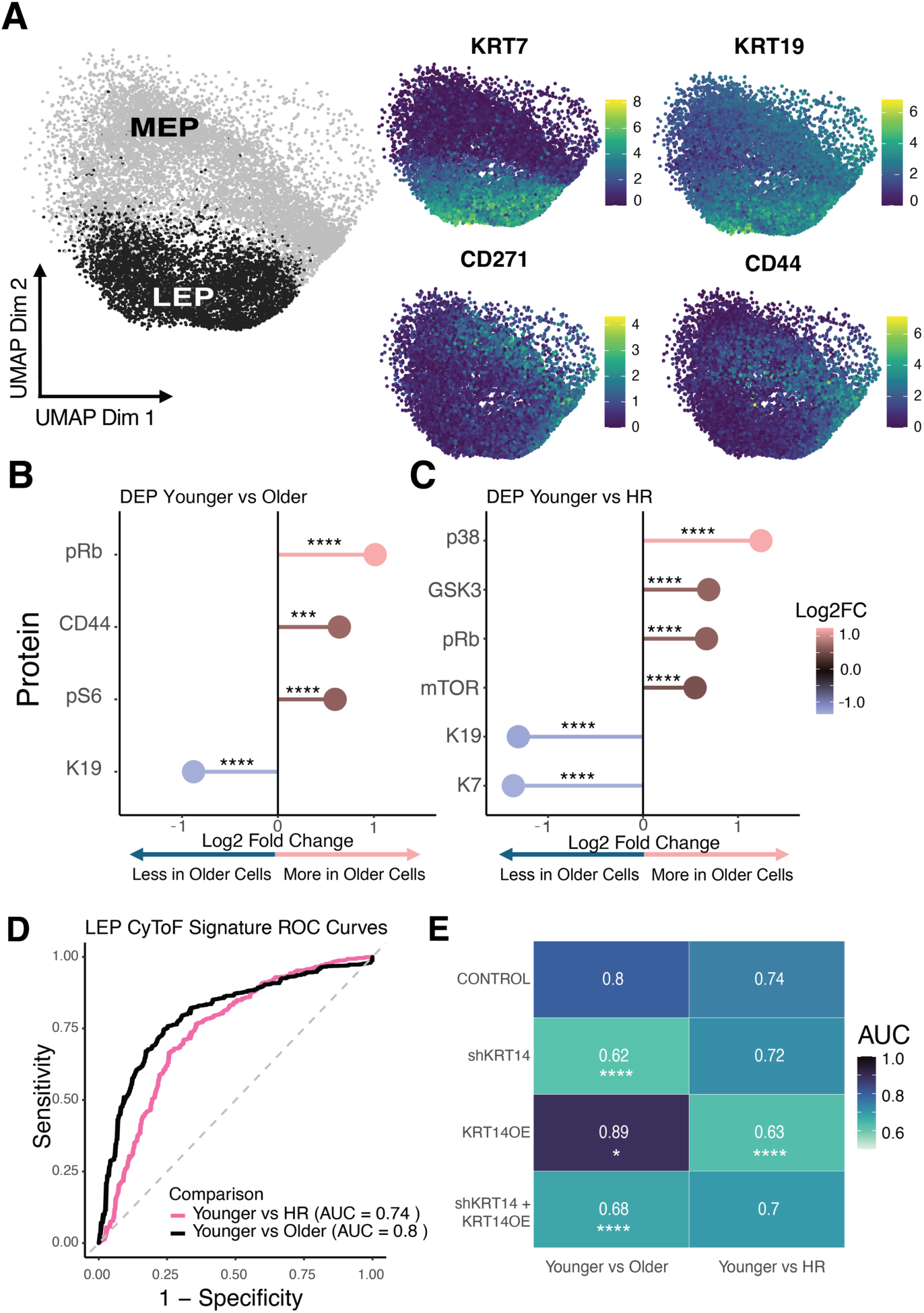
CyTOF Analysis of Luminal Epithelial Cells Across Age and Risk Groups. (A) UMAP visualization of luminal epithelial and myoepithelial cell populations with protein expression mapped for selected markers. (B) Differential protein expression in luminal epithelial cells between younger and older groups and (C) younger vs HR groups. (D) ROC curves generated from GBM models based on differential protein expression, demonstrating predictive capacity for age and high-risk status. (E) Heatmap of AUC values comparing experimental conditions in younger versus older and high-risk groups. Significance was determined using DeLong’s test and compared to control. Dotted line: 95% confidence interval.

Differential protein abundance analysis revealed significant shifts across age and risk groups. Comparisons between younger and older individuals (**Figure 4B**) identified increased expression of pRb, CD44, and pS6 and decreased levels of KRT19 in older cells, which indicated age-associated changes in cell-cycle regulation, signaling, and cytoskeletal integrity. Similarly, comparisons between younger and HR strains (**Figure 4C**) highlighted the altered expression of p38, GSK3, and mTOR.

Beyond control cells, we performed KRT14 overexpression (OE), KRT14 knockdown (shKRT14), and dual modulation (shKRT14 + KRT14OE) experiments to examine how gene expression alterations influence protein signatures in HMECs. The “KRT14OE” construct was designed such that the shKRT14 short hairpin RNA could not target the KRT14 mRNA that was overexpressed, which enabled simultaneous overexpression and knockdown in the same cells and only ablated the autologous KRT14 expression in the double transduction sample. Gradient boosting machine (GBM) models were trained on differentially expressed proteins and produced robust predictive accuracy. GBM algorithms built an ensemble of decision trees sequentially and optimized a loss function through gradient descent. For binary classification tasks such as “younger vs older” or “younger vs high-risk,” GBM minimized the log-loss function, similar to logistic regression, to output probabilities for classification per single cell. Unlike logistic regression, GBM did not assume a linear relationship between input variables and the classification variable, which allowed it to capture nonlinear interactions. ROC curves (**Figure 4D**) demonstrated strong performance in distinguishing younger vs older (AUC = 0.80). This approach highlighted a separation between average risk younger and HR cells (AUC = 0.74). KRT14 OE had a pronounced impact on the age model, which significantly enhanced predictive capacity (AUC = 0.89), while its effects on the HR model were more modest. Similarly, shKRT14 reduced accuracy predominantly in the age model, with a smaller effect on the HR model (**Figure 4E**). However, in myoepithelial cells, which naturally exhibited higher KRT14 expression levels, the impact of KRT14 overexpression on the age model was different. The model’s ability to distinguish between younger and HR cells was less effective in myoepithelial cells as compared to in luminal cells. Myoepithelial cells exhibited a lower AUC of 0.65 for the same comparison, which suggested a weaker separation and implied that high-risk signatures may be more pronounced in luminal epithelial cells (**Figure 5**). Together, these findings suggested that KRT14 modulation caused substantial molecular changes associated with cellular aging, as evidenced by significant shifts in Mechano-RISQ scores. The risk-associated changes in the high-risk model demonstrated that additional pathways are engaged following KRT14 modulation, indicating that age prediction may be influenced by a broader mechanobiological landscape.

**Figure 5.**
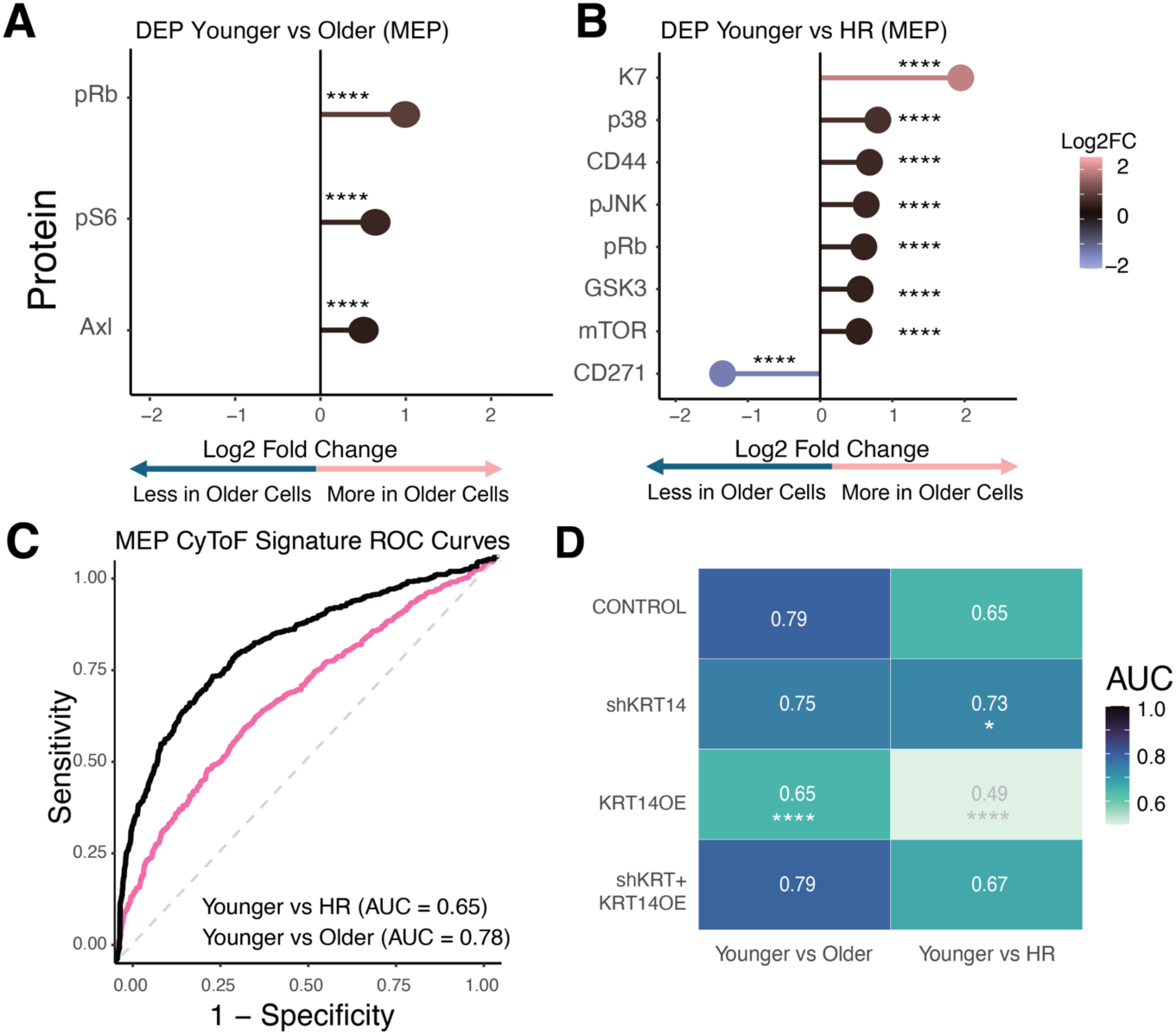
CyTOF Analysis of Myoepithelial Cells Across Age and Risk Groups. (A) Differential protein expression in myoepithelial cells between younger vs older and (B) younger vs HR groups. (C) ROC curves generated from GBM models based on differential protein expression, demonstrating predictive capacity for age and high-risk status. (D) Heatmap of AUC values comparing experimental conditions in younger versus older and high-risk groups. Significance was determined using DeLong’s test and compared to control condition. Significance is indicated as follows: \**P* < 0.05; \*\**P* < 0.01; \*\*\**P* < 0.001, \*\*\*\**P* < 0.0001.

## Discussion

Mechanical phenotyping of single cells offers a pioneering approach to identifying age-associated biologies in HMECs that are relevant to breast cancer risk. High-resolution, single-cell mechanical profiling revealed that aging and high-risk conditions, such as BRCA1/2 mutations and family history of breast cancer, are associated with distinct mechanical states. The ability of MechanoAge to classify cells accurately by age (AUC = 0.95) and to detect deviations in mechanical phenotypes in high-risk individuals suggests that cellular mechanical properties may serve as an early biomarker for breast cancer susceptibility and, thus, as a tool for cancer risk assessment.

The identification of high-risk individuals for breast cancer remains a significant challenge, particularly in cases where genetic mutations are absent or ambiguous. BRCA1 and BRCA2 mutations account for 5-10% of hereditary breast cancer cases; however, the majority of individuals (>90%) diagnosed with breast cancer lack these mutations^21^. Moreover, many fall into intermediate-risk categories, such as those with variants of unknown significance (VUS), a family history of breast cancer without identifiable mutations, or environmental exposome^16^. Traditional risk assessment tools often fail to provide precise or actionable insights for these individuals. Our findings suggest that mechanical profiling may address these challenges by detecting emergent subtle, but highly important, risk-associated cellular changes that are otherwise undetectable. For instance, mechanical changes occur in high-risk groups even in the absence of known genetic mutations, as demonstrated by the elevated Mechano-RISQ scores in both BRCA1/2 mutation carriers and in individuals with a family history of breast cancer. Similarly, younger individuals with a family history of breast cancer exhibited a higher proportion of cells classified as “older,” further supporting the hypothesis that mechanobiological alterations may precede detectable genomic changes that increase cancer susceptibility.

A key strength of this study is the use of mechanical phenotyping for uncovering early indicators of susceptibility in populations that are difficult to stratify with current approaches such as germline DNA sequencing, histopathology, and immunohistochemistry. Mechano-NPS is particularly suited for mechanical phenotyping at scale, given its throughput, which is substantially higher than the gold standards of atomic force microscopy and micropipette aspiration^11,22^. The integration of multidimensional mechanical data in combination with ensemble ML algorithms offers distinct advantages in this context by providing high-resolution, single-cell mechanical profiles in a high-throughput format. Indeed, ML played a critical role in uncovering the combinations of mechanical properties that differentiate otherwise normal HMEC by their age or risk groups. Measurement of mechanical states could serve as a complement to genetic, epigenetic, and lifestyle risk factors to deliver a more comprehensive risk assessment for breast cancer.

Our investigation into the role of the intermediate filament KRT14 highlights its central role in mediating cytoskeletal reorganization during cellular aging. Changes in KRT14 expression in mammary epithelia was among the first overt hallmarks of aging that we observed^8,17^. Overexpression of KRT14 in younger cells induced an “older” mechanical phenotype, and knockdown in older cells attenuated age-associated mechanical changes. These findings indicate that KRT14 is a critical molecular mediator of cytoskeletal remodeling. The effect of KRT14 modulation on mechanical properties corresponds with distinct protein expression changes. GBM models trained on differentially expressed proteins confirmed the importance of KRT14 in distinguishing cells by age (AUC = 0.80) and younger vs high-risk cells (AUC = 0.74). While KRT14 overexpression strongly enhanced age prediction accuracy (AUC = 0.89), its effects on high-risk classification were less pronounced, suggesting that additional pathways contribute to cancer susceptibility beyond KRT14-driven mechanical alterations. The observed changes protein expression resulting from keratin changes align with the hypothesized role of intermediate filaments as signaling scaffolds^23^.

Here, we demonstrated that intermediate filaments are key regulators of these biomechanical profiles. Since all cells contain intermediate filaments, these findings likely extend beyond HMECs, suggesting broader relevance across various cell types and tissues. Age-related biomechanical changes may represent a fundamental hallmark of cellular function, with distinct mechanical phenotypes underlying critical processes in aging, cancer, and potentially other diseases. Recognizing and utilizing these biomechanical markers could greatly enhance early detection, refine risk stratification, and improve targeted intervention strategies. This could be especially significant for those with ambiguous risk profiles, including individuals without known genetic mutations. Critically, incorporating mechanical phenotyping into clinical trials could enhance trial design by allowing earlier, quantifiable endpoints, particularly in prevention studies, which would facilitate a more accurate evaluation of intervention efficacy and risk reduction strategies.

## Material and Methods

### Primary epithelial cells

Primary HMEC strains were derived from tissues discarded from reduction mammoplasties and prophylactic mastectomies (**Figure 1A**) and were propagated in a low-stress medium to passage four, at which point the lineage heterogeneity of the original mammary epithelium is maintained. Primary HMEC strains were maintained as described previously ^13,24^. The HMEC strains used in this study are listed in **Table S1**. All cultures were screened regularly and remained mycoplasma-free throughout the study. For lineage-specific mechanical measurements, cells were separated using Stem Cell’s EasySep technology and stained with CD271 and CD133 antibodies. For transduction studies, HMECs were transduced with lentiviral particles in the presence of polybrene (10 µg/ml) and were selected using puromycin (2 µg/mL) or hygromycin (10 µg/mL).

### Mechano-NPS Device Design

The mechano-NPS devices used in this study consisted of a 22.3 µm-high polydimethylsiloxane (PDMS) microfluidic channel bonded to a glass substrate with pre-defined platinum (Pt) and gold (Au) contact pads. The channel consists of a series of nodes (n) and pores (p) on either side of a single “contraction channel”. Each node is 50 µm x 85 µm (L x W), each pore is 700 µm x 22 µm (L x W), and the contraction channel is 3000 µm x 10.5 µm (L x W). The contraction channel width, *w*_c_, was chosen to apply an average strain, *ε* = (*D*_cell_-*w*_c_)/*D*_cell_ ∼0.3, over a sufficient amount of time to each cell transiting the microfluidic device. Here, *D*_cell_ is the free cell diameter. The recovery length segment is 700 um long.

### Mechano-NPS Device Fabrication

Mechano-NPS devices were fabricated, as previously published^11,22,25^. Briefly, to create the PDMS microfluidic molds, standard photolithography was first used to fabricate negative-relief masters on polished silicon wafers. PDMS (Sylgard 184, Dow Corning), mixed at a ratio of 10:1 pre-polymer: curing agent and degassed, was then poured onto the negative-relief masters and cured at 85°C for 2 hrs. A slab of PDMS with the embedded microfluidic channel was excised from the master, and input and outlet ports were cored with a 1.5 mm diameter biopsy punch. Completion of the mechano-NPS device involved exposing a PDMS mold and a glass substrate with pre-defined electrodes to an oxygen plasma (Harrick Plasma, 450 mTorr (60 Pa), 30 W, 2 min) and aligning and mating the two together. Permanent bonding of the glass substrate and PDMS mold was accomplished by heating the device on a hotplate at 85°C for 2 hrs. To fabricate the Pt electrodes and Au contact pads onto the glass substrates, standard photolithography was used for patterning. A trilayer of thin metal film (50/250/250 Å titanium/Pt/Au) was deposited via electron-gun evaporation and a subsequent lift-off with acetone was performed. Gold wet etch (Gold Etchant TFA, Transene Company) was used to expose the Pt electrodes.

### Mechano-NPS Device Measurement

Single-cell suspensions of HMECs (∼2 x 10^5^ cells/ml) were resuspended in PBS and immediately injected into the device. A constant DC voltage (1V) was applied across the channel, and a four-terminal measurement of the current was used to measure the modulated current pulses caused cells transiting the channel when a pressure of 25 kPA was used. We employed custom-written software (see Zenodo 10.5281/zenodo.14927530) to extract both the magnitude and duration of each current sub-pulse (Δ*I*_np_, Δ*I*_c_, Δ*T*_cont_, and Δ*T*_r_). Δ*I*_np_ corresponds to the current drop as the cell enters the node-pore region at the beginning of the device, μ*I*_c_ is the current drop as the cell enters the contraction channel, Δ*T*_cont_ is the cell’s transit time through the contraction channel, and μ*T*_r_ is the time necessary for the cell to relax to its original shape after deformation. If a cell remains deformed past the observation period, its relaxation time is recorded as “Inf”.

### Determination of the Free Cell Diameter, *D*_cell_, Transverse Deformation, Recovery Time, and Whole Cell Deformability Index, *wCDI*

The magnitude of the current sub-pulse, Δ*I_np_*/*I*, produced in the node-pores prior to the contraction channel provides information on the *D*_cell_. Specifically,

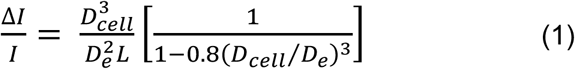

where *D*_e_ and *L* is the effective diameter and length of the channel, respectively. See Ref.^11^ for more details on the derivation of Eq. (1). *D*_e_ is determined by measuring polystyrene microspheres of known size with the microfluidic channel. Once *D*_e_ is known, then *D*_cell_ of a screened cell can be numerically solved in Eq. (1) using the obtained values of Δ*I*_np_/*I*.

To determine a cell’s transverse deformation^11^, we assume that the cell is an oblate spheroid when in the contraction channel. Its volume is *V*_deform_ = ρχ*w*_c_(*L*_deform_)^2^/6 where *L*_deform_ is the cell’s deformation length. As the magnitude of the current change in the contraction channel, μ*I*_c_/*I* is proportional to the volume ratio of the deformed cell and contraction channel, *V*_deform_/*V*_contraction_, the transverse deformation of the cell is thus *ο*_deform_ = *L*_deform_ /*D*_cell_.

To determine the recovery time of a cell after applied strain, μ*T*_r_, we note the time required for the sub-pulses produced by the cell *after* exiting the contraction channel to return to the same shape and magnitude as those produced by the cell *prior* to entering the contraction channel. For the device dimensions used in this study, the temporal window is ∼75 ms for the applied flow rate.

The *wCDI* is a dimensionless parameter, which was previously shown to be inversely related to cortical tension or Young’s modulus^11^. This parameter is defined as,

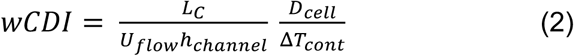

where *U*_flow_ is the fluid velocity in the node section leading into the contraction channel, *L*_c_ is the length of the contraction channel, and *h*_channel_ is the contraction-channel height. *U*_flow_, *L*_c_, and *h*_channel_ are fixed values for any given experiment, and consequently, *D*_cell_ and ΔT_cont_ are the key parameters in the wCDI.

### Classifying age groups using mechano-NPS data

A database of mechano-NPS measurements was created from HMECs of 18 women, including 1381 cells in the training set and 661 cells in the validation dataset. Cells were manually classified based on chronological age into two groups: “older” (>55 years of age) and “younger” (<35 years of age). The modeling was performed in the R programming environment using the caret package. A detailed pseudocode representation of this workflow is included in **Algorithm 1**. Variables were derived from the non-continuous “RecoveryTime” variable. Dummy variables were created to represent five distinct recovery time intervals. A Yeo-Johnson transformation was applied to improve the normality of the data distribution and ensure variance homogeneity. The data were centered and scaled after the transformation to standardize the features. Three ML algorithms were employed to model the relationship between predictor variables and age categories: Bagged Decision Trees^26^, Random Forest^27^, and Extremely Randomized Trees^28^, all implemented using the caret package^29^. Hyperparameter tuning was conducted with 30 different values to optimize model performance concerning the area under the receiver operating characteristic curve (ROC AUC). Model training was performed using repeated k-fold cross-validation (10 folds, five repeats) to ensure robust performance estimation optimized for ROC AUC. A down-sampling strategy was employed to address minor class imbalance in the outcome variable. An ensemble model was constructed using the caretStack function from the caretEnsemble package^30^ to enhance predictive performance. The ensemble combined the predictions of the three individual models using a Generalized Boosted Regression Model (GBM)^31^ as the meta-learner. The GBM was configured with a tuning length of 10, and model training was performed using 5-fold cross-validation. The performance metric was set to ROC AUC, and a down-sampling strategy was maintained to address class imbalance. The final optimized ensemble model was applied to the validation dataset to generate class predictions and associated probabilities.

### Mechano-Risk Index for Single-cell Quantification (RISQ) Score Calculation

The Mechano-RISQ score quantifies the deviation of cell classification as “older” relative to a defined baseline error rate in the context of mechanical and age-related phenotype predictions. The baseline error rate represents the proportion of cells misclassified as “older” in the core model, determined from average risk HMECs. This value was used as a reference to calculate the relative increase or decrease in the proportion of misclassified cells across experimental conditions. To calculate the Mechano-RISQ score for each experimental condition, the percentage of misclassified cells was compared to the baseline error rate to compute the ratio of deviation. A ratio of 1.0 indicates no deviation from the baseline, while values above or below 1.0 reflect an increase or decrease in the misclassification rate, respectively.

### CyTOF

Pre-conjugated antibodies were obtained from Standard Biotools (San Francisco, USA) (**Supplementary Table 2**). Unlabeled antibodies were conjugated in-house using the Maxpar® X8 Antibody Labeling Kit (Standard BioTools) or the MIBItag Conjugation Kit (Ionpath, Menlo Park, CA, USA). Primary HMECs were stained for viability with cisplatin according to the manufacturer’s protocol and then fixed with paraformaldehyde (PFA). Samples were incubated in PBS before primary barcoding into three separate pools using the. After barcoding, the combined pool was aliquoted for titration, validation, and acquisition. Titration samples were used to optimize marker concentrations, validation samples confirmed titration accuracy, and acquisition samples were reserved for final data collection. The combined barcode pool washed and stained with a mixture of metal-conjugated antibodies against surface proteins (**Supplementary Table 2**). After staining, cells were washed in CSB, fixed in 2% PFA, and permeabilized in methanol. Subsequently, cells were washed in CSB and stained with a cocktail of metal-conjugated antibodies for intracellular proteins (**Supplementary Table 2**). Following staining, cells were fixed in 2% PFA with Cell-ID™ Intercalator-Ir (Standard BioTools). For acquisition cells were diluted to was acquired using the tuned Helios mass cytometer with the Super Sampler. Differentially expressed proteins (DEPs) between groups were analyzed on a per-protein basis, applying a non-parametric Wilcoxon rank-sum test to detect significant differences in expression. Multiple hypothesis testing corrections were applied using the Bonferroni method, adjusting P-values. Proteins were reported as significant if they met predefined thresholds of adjusted P-value < 0.05 and |LFC| > 0.5.

### CyTOF Expression Signatures

CyTOF differential protein signatures were modeled with gradient-boosting machines. A stratified 70/30 train–test split and five-fold cross-validation maximized AUC, and test-set ROC curves (sensitivity vs 1-specificity) summarized single-cell age- and risk-classification performance for direct model comparisons.

### Power Analyses

Power analyses remain an active area of research in machine learning, particularly in emerging applications like ours that explore the mechanical properties of single cells in the aging context. Given the speculative nature of applying traditional power analysis methods to these novel settings, we conducted a learning curve analysis to guide our experimental design regarding the training database size (**Supplemental Figure 2**). Our analysis—based on random sampling repeated five times across sample sizes ranging from 40 to 1300—reveals an increase in ROC AUC from approximately 0.68 to 0.87. The first derivative of ROC AUC with respect to sample size (*d(AUC)/d(n)*) trends toward zero, indicating that the marginal performance improvement diminishes as the sample size increases. Additionally, the AUC estimates’ standard error (SE) decreases with larger sample sizes, reflecting enhanced stability and reliability of the performance metric. In machine learning, such a plateau in the learning curve combined with reduced variability suggests that the model is approaching its optimal performance regime, implying that further data collection would likely yield diminishing returns relative to its cost.

To assess the sensitivity of Crawford–Howell test, we conducted a simulation-based power analysis. Specifically, we simulated 10,000 iterations of a control sample (n = 18) drawn from a normal distribution with a mean of 1.2 and a standard deviation of 1, parameters derived from our normative dataset.

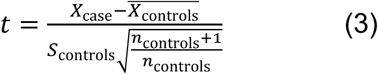

A single-case value was generated for each iteration corresponding to an effect size (Cohen’s d) of 3. The Crawford–Howell t statistic was calculated as Eq (3). A two-tailed test at α=0.05 was applied, and power was estimated to be above 0.97.

## Acknowledgments

We gratefully acknowledge our patient advocates, Susan Samson and Sandy Preto, for providing much-needed context. This research was partially supported from NIH R01CA237602, U01CA244109, and BC181737(MAL), 1R01EB024989 (LS and ML), the American Cancer Society – Fred Ross Desert Spirit Postdoctoral Fellowship (PF-21-184-01-CSM) (SH), a University of Bergen PhD fellowship to (SMG), a Siebel Scholars Foundation Fellowship (KLC), a UC Berkeley Mentored Research Award (TT), and a Chancellor’s Fellowship (CC). The research reported in this publication included work performed in the Integrative Genomics and Analytical Cytometry Cores supported by the National Cancer Institute (NCI) of the National Institutes of Health under award number P30CA033572. The content is solely the responsibility of the authors and does not necessarily represent the official views of the National Institutes of Health. Mass cytometry was performed at the Flow & Mass Cytometry Core Facility, Department of Clinical Science, University of Bergen. The Helios mass cytometer was funded by Trond Mohn Research Foundation. Figure 1 A was created in BioRender. Hinz, S. (2025) https://BioRender.com/u93m128.

## Author Contributions

Conceptualization: SH, MM, LLS, and MAL,

Writing – Original Draft: SH and MAL.

Writing – Review & Editing: SH, MM, SMG, LDY, LLS, JBL, and MAL.

Investigation: SH, SMG, MM, and JCL.

Device Fabrication: KLC, TT, CC, and EJH.

Formal Analysis: SH.

Data Curation: SH and SMG. Visualization: SH.

Biospecimen Provision: VES.

## Data and materials availability

Available upon request, subject to a Material Transfer Agreement (MTA). CyTOF data is deposited through 10.5281/zenodo.15004690.

## Supplementary Materials

Supplemental Figures

Supplemental Tables

---

**Algorithm 1:**
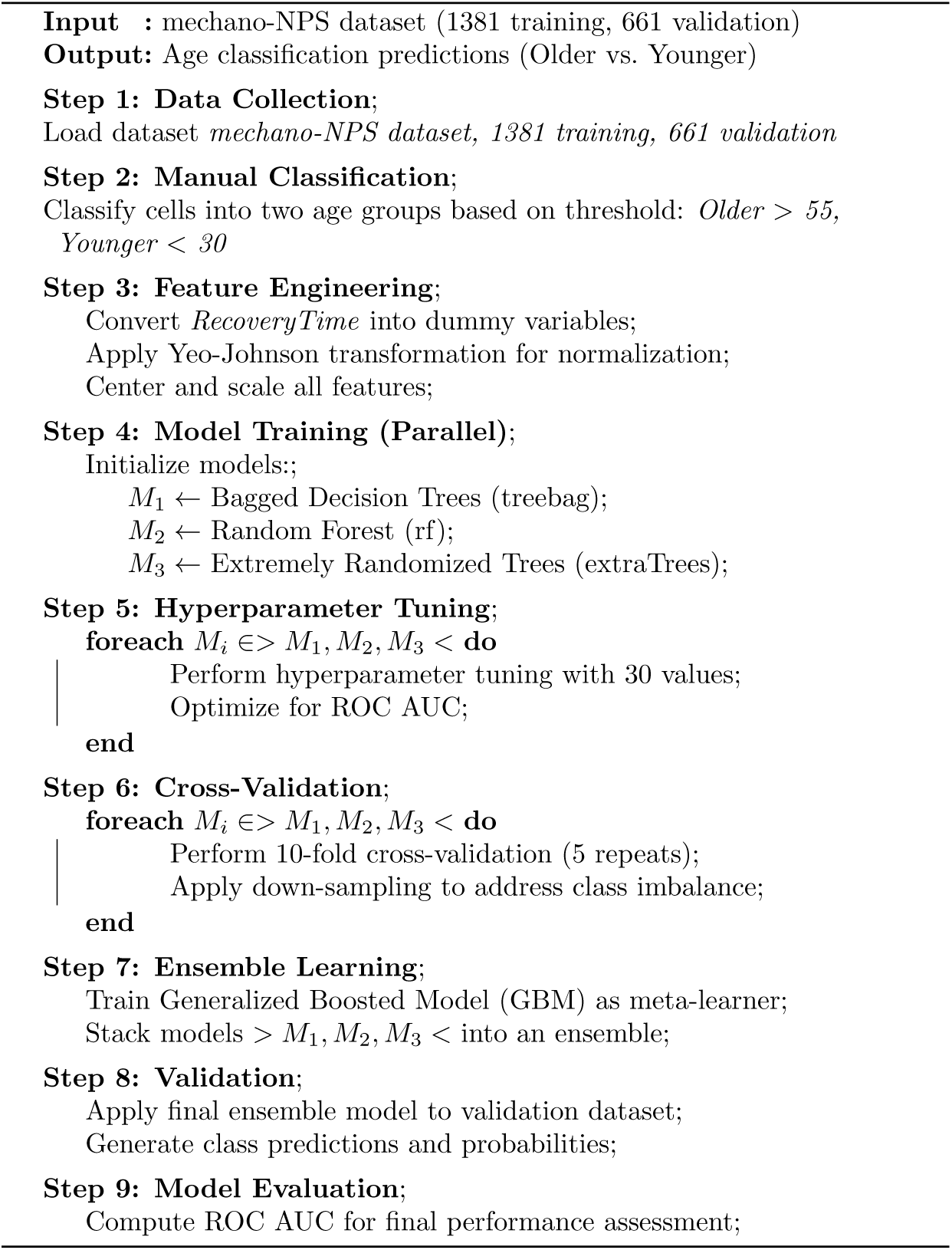
Machine Learning Pipeline for Age Classification Pseudocode for MechanoAge

## Supplemental Figures

**Supplemental Figure 1:**
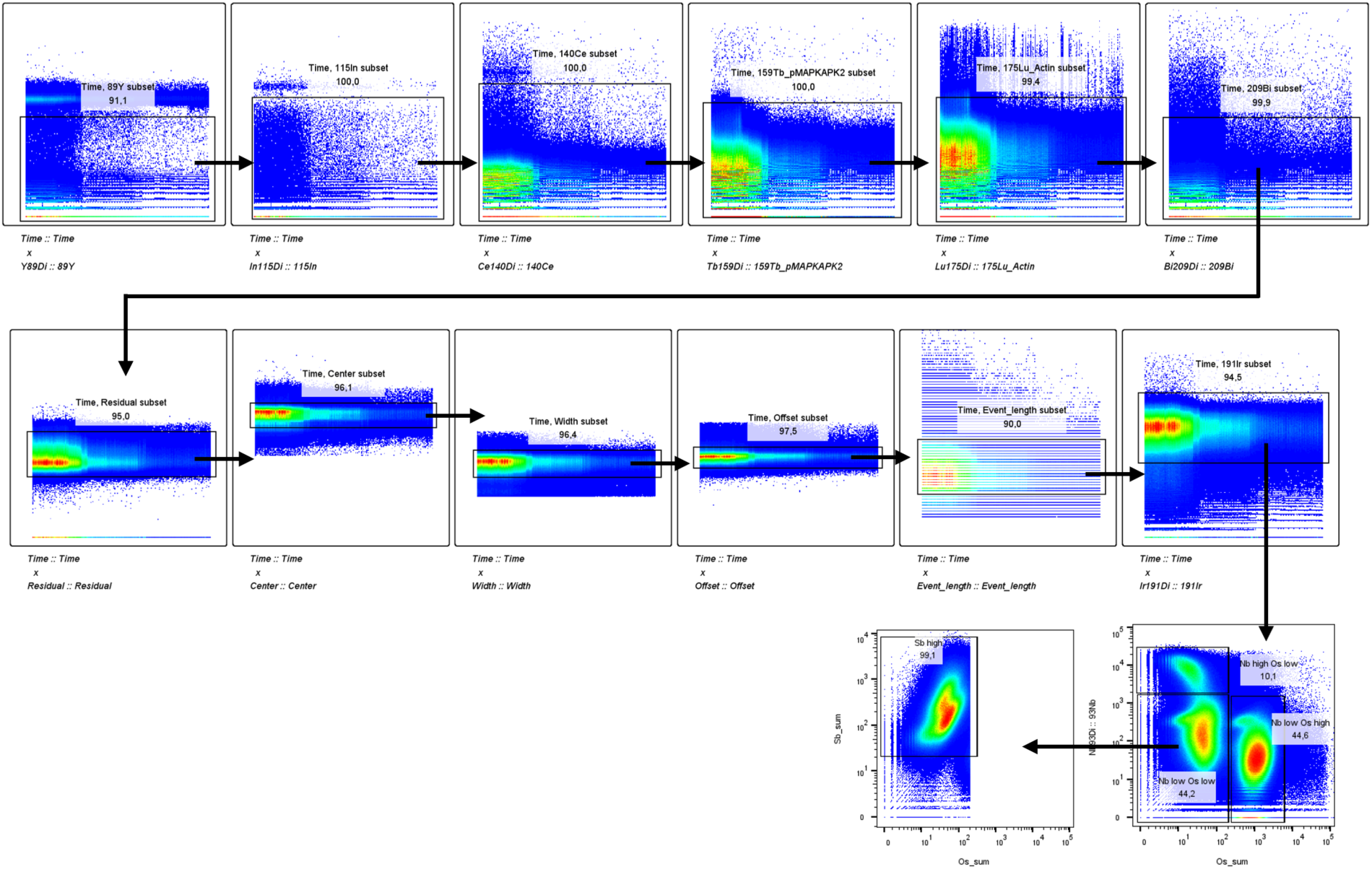
Gating strategy for mass cytometry data. Gating was performed on normalized and concatenated events. Top row of gates gate on beads for every channel. Second row applies gating on gaussian parameters and DNA (191Ir). Third row gates out barcode pools based on staining of niobium, osmium, and antimony.

**Supplemental Figure 2:**
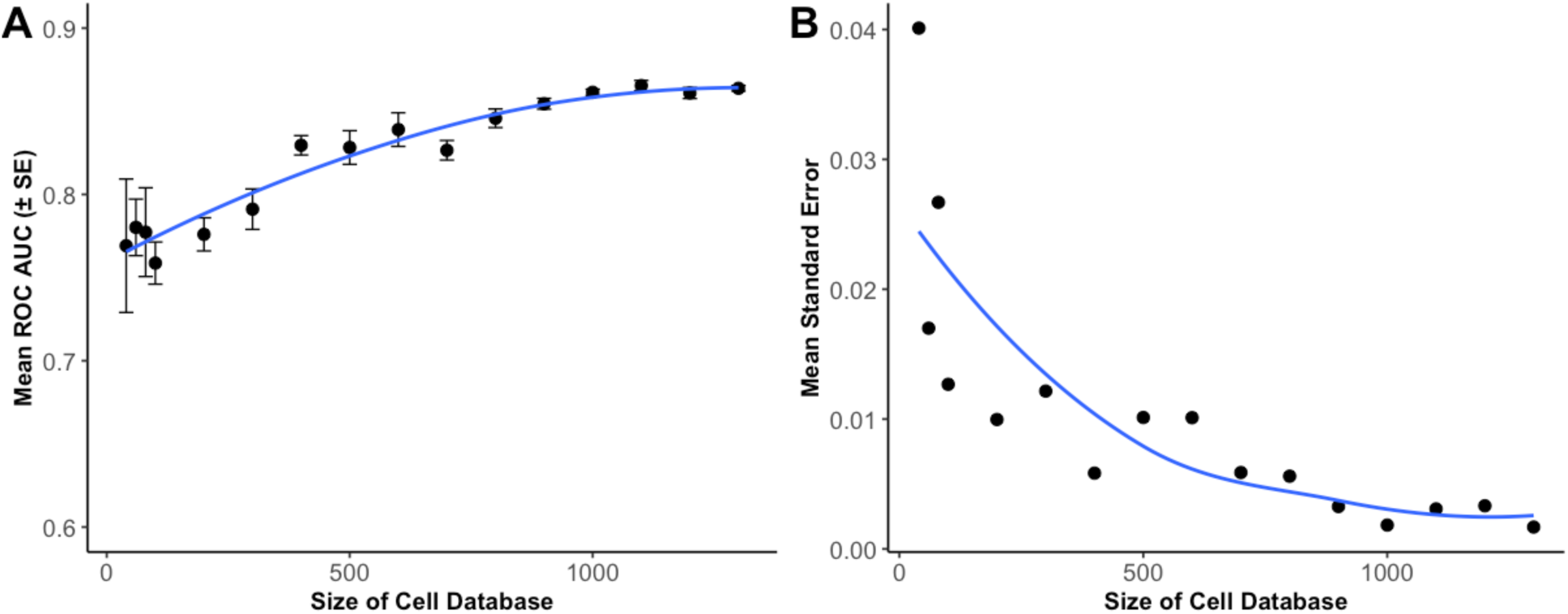
Learning Curve Analysis for Estimating Cell Database Size Requirements. (A) Mean ROC AUC (± SE) plotted against the size of the cell database. (B) Mean standard error of the AUC for the same samples.

## Supplemental Tables

**Supplementary Table 1:**
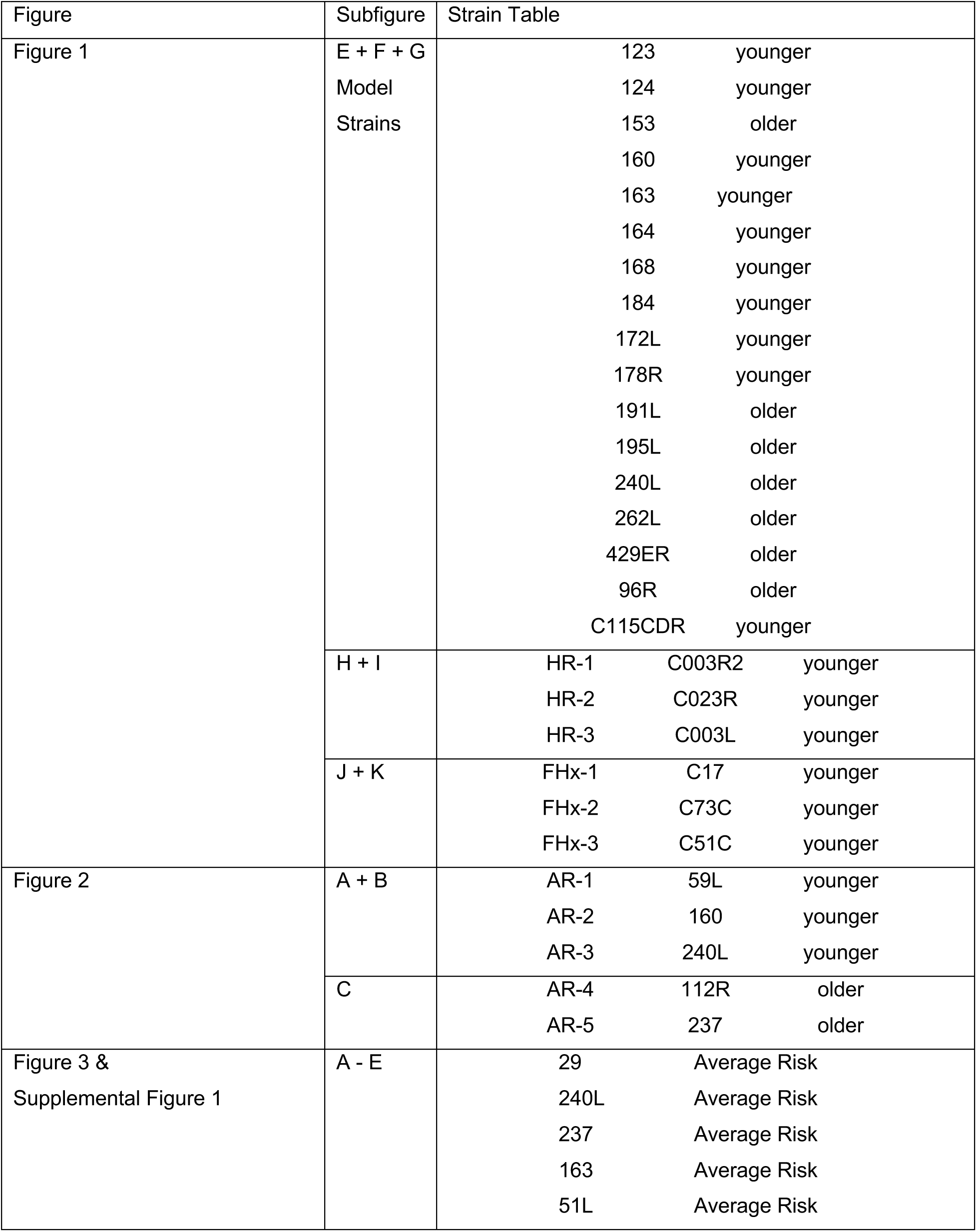
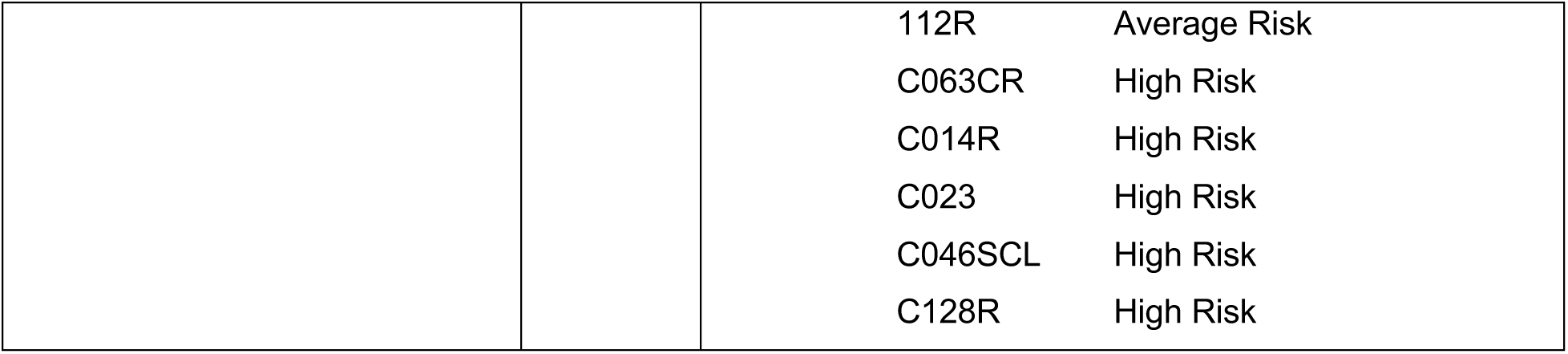
Specimen Table.

| Figure | Subfigure | Strain Table |  |  |
| --- | --- | --- | --- | --- |
| Figure 1 | E + F + G<br>Model<br>Strains | 123 | younger |  |
|  |  | 124 | younger |  |
|  |  | 153 | older |  |
|  |  | 160 | younger |  |
|  |  | 163 | younger |  |
|  |  | 164 | younger |  |
|  |  | 168 | younger |  |
|  |  | 184 | younger |  |
|  |  | 172L | younger |  |
|  |  | 178R | younger |  |
|  |  | 191L | older |  |
|  |  | 195L | older |  |
|  |  | 240L | older |  |
|  |  | 262L | older |  |
|  |  | 429ER | older |  |
|  |  | 96R | older |  |
|  |  | C115CDR | younger |  |
|  | H + I | HR-1 | C003R2 | younger |
|  |  | HR-2 | C023R | younger |
|  |  | HR-3 | C003L | younger |
|  | J + K | FHx-1 | C17 | younger |
|  |  | FHx-2 | C73C | younger |
|  |  | FHx-3 | C51C | younger |
| Figure 2 | A + B | AR-1 | 59L | younger |
|  |  | AR-2 | 160 | younger |
|  |  | AR-3 | 240L | younger |
|  | C | AR-4 | 112R | older |
|  |  | AR-5 | 237 | older |
| Figure 3 &<br>Supplemental Figure 1 | A - E | 29 | Average Risk |  |
|  |  | 240L | Average Risk |  |
|  |  | 237 | Average Risk |  |
|  |  | 163 | Average Risk |  |
|  |  | 51L | Average Risk |  |

|  |  |  |  |
| --- | --- | --- | --- |
|  |  | 112R | Average Risk |
|  |  | C063CR | High Risk |
|  |  | C014R | High Risk |
|  |  | C023 | High Risk |
|  |  | C046SCL | High Risk |
|  |  | C128R | High Risk |

**Supplementary Table 2:** Antibodies for mass cytometry.

| Element | Mass | Target | staining location | Host | Clone | Vendor | Cat# | Lot# |
| --- | --- | --- | --- | --- | --- | --- | --- | --- |
| Nd | 145 | MEK 1/2 | Intracellular | Rabbit | MEK12S<br>217S221<br>-H2 | Thermo<br>Fisher | MA5-28037 | XC3547831 |
| Nd | 146 | mTOR | Intracellular | Mouse | A17024A | BioLegend | 610302 | B273519 |
| Sm | 147 | pSTAT5 | Intracellular | Mouse | 47 | Fluidigm | 3147012<br>A | 1701901 |
| Nd | 148 | K19 | Intracellular | Mouse | BA17 | Thermo<br>Fisher | 14-9898-82 |  |
| Sm | 149 | CD271 | Surface | Mouse | C40-1457 | Fluidigm | 3149017<br>B | 2181507 |
| Nd | 150 | pRb | Intracellular | Mouse | J112-906 | Fluidigm | 3150013<br>A | 0361504 |
| Sm | 152 | pAkt | Intracellular | Rabbit | D9E | Fluidigm | 3152005<br>A | 2103273-09 |
| Eu | 153 | pSTAT1 | Intracellular | Mouse | 4a | Fluidigm | 3153005<br>A | 1501704 |
| Sm | 154 | Vimentin | Intracellular | Rabbit | D21H3 | Fluidigm | 3154014<br>A | 0591704 |
| Gd | 155 | GSK3 | Intracellular | Rabbit | 19H1L12 | Thermo<br>Fisher | 702230 | 2084110 |
| Gd | 156 | p38 | Intracellular | Rabbit | D3F9 | Fluidigm | 3156002<br>A | 2631813 |
| Gd | 157 | Axl | Surface | Mouse | 108724 | R&D | MAB154 | GCV0821041 |
| Gd | 158 | pSTAT3 | Intracellular | Mouse | 4/P-STAT3 | Fluidigm | 3158005A | 0132015 |
| Tb | 159 | pMAPKAPK2 | Intracellular | Rabbit | 27B7 | Fluidigm | 3159010A | 1501813 |
| Gd | 160 | CD133 | Surface | Mouse | AC133 | Miltenyi Biotec |  | 130-090-422 |
| Dy | 161 | Ki-67 | Intracellular | Mouse | B56 | Fluidigm | 3161007B | 0641801 |
| Dy | 162 | MST2 | Intracellular |  | Polyclonal | LSBio |  |  |
| Dy | 163 | pPEAK | Intracellular | Rabbit | Polyclonal | Milipore | ABT52 | 3439693 |
| Dy | 164 | K7 | Intracellular | Mouse | RCK105 | Fluidigm | 3164028D | 2112418-15 |
| Er | 166 | NF-kBp65 | Intracellular | Mouse | K10-895.12.50 | Fluidigm | 3166006A | 1111703 |
| Er | 167 | YAP1 | Intracellular | Mouse | H-9 | santa cruz biotech | sc-271134 |  |
| Er | 168 | pSTAT6 | Intracellular | Mouse |  | Fluidigm | 3168012A | 1422005 |
| Yb | 171 | pERK 1/2 | Intracellular | Rabbit | D13.14.4E | Fluidigm | 3171010A | 0762004 |
| Yb | 172 | pS6 | Intracellular | Mouse | N7-548 | Fluidigm | 3172008A | 2104622-07 |
| Yb | 173 | CD44 | Surface | Rat | IM7 | Fluidigm | 3150018B |  |
| Yb | 174 | pJNK | Intracellular | Rabbit | Polyclonal | R&D | AF1205 | HCT0620111 |
| Yb | 176 | c-Myc | Intracellular | Mouse | 9E10 | Fluidigm | 3176012B | 1841816 |

